# Irreversible inactivation of lactate racemase by sodium borohydride reveals reactivity of the nickel-pincer nucleotide cofactor

**DOI:** 10.1101/2022.11.09.515893

**Authors:** Santhosh Gatreddi, Dexin Sui, Robert P. Hausinger, Jian Hu

## Abstract

The nickel-pincer nucleotide (NPN) cofactor discovered in lactate racemase from *Lactiplantibacillus plantarum* (LarA_*Lp*_) is essential for the activities of racemases/epimerases in the highly diverse LarA superfamily. Prior mechanistic studies have established a proton-coupled hydride-transfer mechanism for LarA_*Lp*_, but direct evidence showing that hydride attacks the C4 atom in the pyridinium ring of NPN has been lacking. Here, we show that sodium borohydride (NaBH_4_) irreversibly inactivates LarA_*Lp*_ accompanied by a rapid color change of the enzyme. The drastically altered ultraviolet-visible spectra during NaBH_4_ titration supported hydride transfer to C4 of NPN, and the concomitant Ni loss unraveled by mass spectrometry experiments accounted for the mechanism-based inactivation. High resolution structures of LarA_*Lp*_ revealed a substantially weakened C-Ni bond in the metastable sulfite-NPN adduct where the NPN cofactor is in the reduced state. These findings allowed us to propose a mechanism of LarA_*Lp*_ inactivation by NaBH_4_ that provides key insights into the enzyme-catalyzed reaction and sheds light on the reactivity of small molecule NPN mimetics.

## Introduction

LarA from *Lactiplantibacillus plantarum* (LarA_*Lp*_) is a lactate racemase that interconverts the L- and D-enantiomers of the α-hydroxyacid.^1, 2^ As the ninth nickel-dependent enzyme discovered in nature, LarA_*Lp*_ utilizes a novel nickel-pincer nucleotide (NPN) cofactor (**Scheme 1A**)^3^ to catalyze the isomerization reaction.^4^ The coenzyme is synthesized from nicotinic acid adenine dinucleotide by consecutive actions of the LarB carboxylase/hydrolase,^5^ LarE sulfur transferase,^6^ and LarC nickel insertase.^7^ The most prominent characteristic of the NPN cofactor is the direct bonding between Ni^2+^ and the C4 atom in its pyridinium ring, representing the first C-Ni bond (other than transient intermediates) identified in a biological system. Since the discovery of the NPN cofactor in 2015, the large LarA superfamily, previously denoted as the DUF2088 family, was shown to consist of highly diverse family members catalyzing racemization/epimerization reactions on a variety of α-hydroxyacids with many representatives catalyzing unknown reactions.^8^

The reaction mechanism of LarA_*Lp*_, the founding member of the LarA superfamily, has been extensively studied.^9^ A proton-coupled hydride-transfer (PCHT) mechanism (**Scheme 1B**), where each half reaction is utilized in D-/L-lactate dehydrogenases,^1^ has been proposed based on the enzyme structure, a substrate kinetic isotope effect, identification of the pyruvate intermediate, spectroscopic changes with added substrate, and computational investigations.^9, 10, 11^ Supporting the proposal for hydride transfer to C4 of the NPN cofactor, the addition of sulfite to the enzyme led to spectroscopic changes that were associated with formation of a sulfite-NPN covalent adduct which was revealed in a crystal structure of LarA_*Lp*_.^9^ Nevertheless, direct evidence of hydride transfer to C4 of NPN is still lacking.

In this work, we provide compelling biochemical, spectroscopic, and structural evidence supporting the hydride reactivity of C4 in the NPN cofactor. LarA_*Lp*_ was found to be irreversibly inactivated by sodium borohydride (NaBH_4_) accompanied by a rapid color change of the enzyme. Ultraviolet-visible (UV-Vis) spectroscopy, used to monitor the titration of LarA_*Lp*_ with NaBH_4_, indicated that the hydride is transferred to C4 of NPN, and mass spectrometry (MS) experiments revealed an unexpected Ni loss caused by hydride attack and further reactivity involving C4. High-resolution structures of LarA_*Lp*_ revealed a weakened C-Ni bond in the sulfite-NPN adduct, where the NPN cofactor is in the reduced state, providing a structural basis of LarA_*Lp*_ inactivation by NaBH_4_. The potential role of the pyruvate reaction intermediate in protecting the vulnerable reaction center from permanent inactivation is discussed. Overall, these new findings provide key insights into the PCHT mechanism of LarA_*Lp*_.

## Results and Discussion

### NaBH_4_ irreversibly inactivates LarA_Lp_

Hydride transfer is the key step of the proposed mechanism of LarA_*Lp*_,^9^ leading us to wonder what would happen if the enzyme was treated with NaBH_4_, a hydride precursor commonly used in protein chemistry. As shown in **Figure 1A**, treatment of the purified LarA_*Lp*_ with a freshly prepared NaBH_4_ solution at neutral pH led to a dose-dependent inactivation. Removing the residual NaBH_4_ by ultrafiltration did not lead to recovery of the activity (**Figure 1B**), indicating that the inhibition is irreversible.

**Figure 1.**
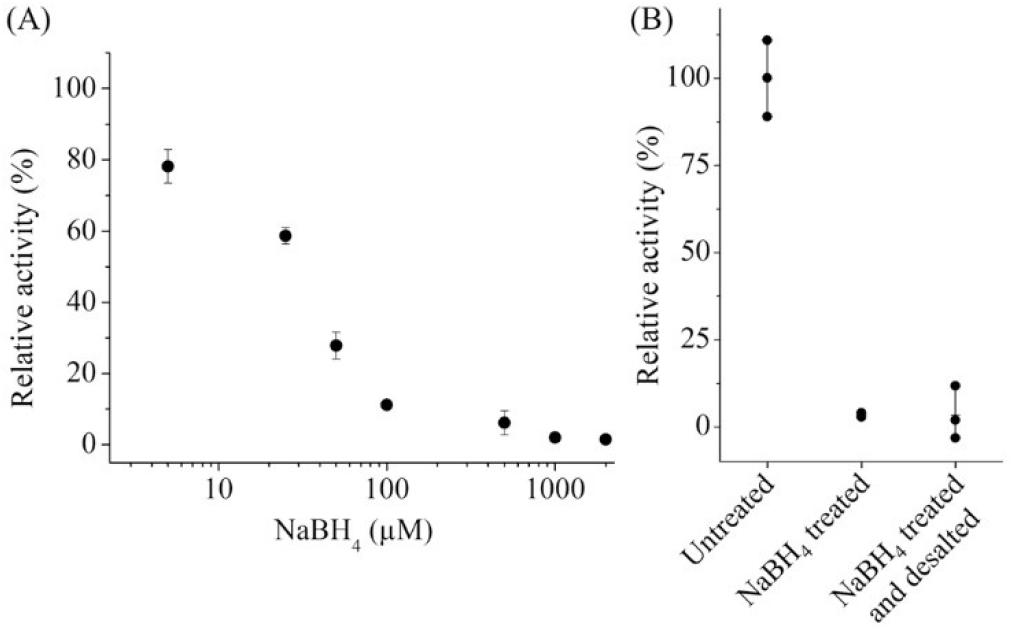
Irreversible inactivation of LarA_*Lp*_ by NaBH_4_. (**A**) Dose-dependent inactivation of LarA_*Lp*_ by a 30-min incubation with the indicated concentrations of NaBH_4_. The purified enzyme (∼20 μM) was treated with freshly prepared NaBH_4_ in buffer containing 50 mM Tris-HCl, pH 7.5, 125 mM NaCl, and ∼0.1 μM Na_2_SO_3_ at room temperature. The activity of NaBH_4_ treated enzyme is expressed as the percentage of the activity of the untreated enzyme. (**B**) Attempt to rescue the activity of LarA_*Lp*_ that had been treated with 2 mM NaBH_4_ by removing residual reductant through desalting. In both (A) and (B), three replicates were included for each condition. The error bars indicate ±S.D.

### UV-Vis spectroscopic characterization of LarA_Lp_ treated with NaBH_4_

The addition of NaBH_4_ to LarA_*Lp*_ that was co-purified with sulfite, but not to the buffer used during enzyme purification, led to a rapid color change from light brown to yellow within seconds (**Figure S1**). Sulfite is routinely added during purification of LarA_*Lp*_ as an activity protectant,^3^ and we have shown previously that sulfite can form a covalent adduct with the NPN cofactor.^9^ We monitored the spectral changes of LarA_*Lp*_ with bound sulfite during a NaBH_4_ titration experiment (**Figure 2**). Consistent with a previous report, untreated LarA_*Lp*_ in the presence of sulfite exhibited a stable yellow color with two major absorption peaks at 380 nm and 440 nm along with a broad minor peak at 540 nm and a shoulder peak at 320 nm.^9^ As shown in **Figure 2A**, the addition of NaBH_4_ drastically changed the spectrum, indicating that the sulfite-NPN cofactor and/or the local environment of the chromophore has been changed by NaBH_4_. Specifically, the 380 nm and 540 nm peaks were significantly suppressed, while the peak at 440 nm increased in intensity with a 10 nm blue shift. The decrease in the 380 nm/440 nm ratio suggested that the sulfite-NPN adduct may have decomposed.^9^ In addition, the peak at 320 nm, which is an indicator of the S-C bond formation between sulfite and NPN (or NAD^+^) at C4, also was diminished. Notably, an increased absorption at 340 nm was apparent in the difference UV-Vis spectra (**Figure 2B**). Because an absorption at 340 nm is a hallmark of hydride transfer to NAD^+^, the 340 nm transition associated with NaBH_4_ titration is a strong indicator of hydride transfer to C4 of the NPN cofactor. Consistent with this interpretation, adding lactate to the sulfite-free LarA_*Lp*_ led to a similar spectral change at 340 nm.^9^ Further analysis of the difference spectra showed that the changes at the mentioned wavelengths have similar dose-dependence profiles, suggesting that the spectral changes are primarily associated with a single process during the titration (**Figure S2**). Collectively, the data suggested a scenario where the sulfite-NPN adduct formed during sample preparation decomposed as a hydride derived from NaBH_4_ attacked C4 of the NPN cofactor.

**Figure 2.**
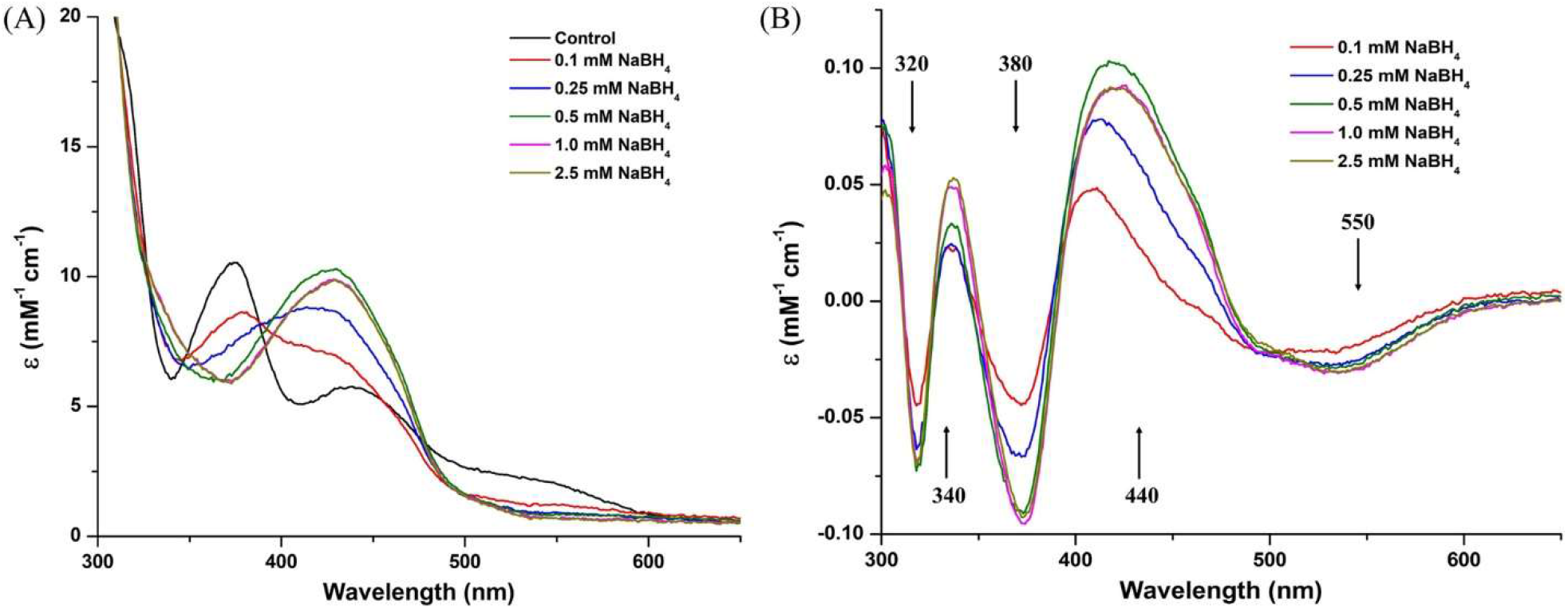
UV-Vis spectroscopic changes of LarA_*Lp*_ that was titrated with NaBH_4_. The monomeric LarA_*Lp*_ (∼20 μM) collected by size-exclusion chromatography (50 mM Tris-HCl, pH 7.5, 125 mM NaCl, and 50 μM Na_2_SO_3_) was treated with NaBH_4_ (0.1-2.5 mM) for 10-15 min at room temperature. The spectra were recorded after bubbles (hydrogen gas) dissipated. (**A**) Superimposed UV-Vis spectra of LarA_*Lp*_ after incubation with NaBH_4_ at the indicated concentrations. (**B**) UV-Vis difference spectra. The difference spectra were generated by subtracting the spectrum of the control sample (untreated) from the spectra of the panel A samples treated with NaBH_4_. The directions of the arrows associated with the indicated wavelengths indicate the direction of change for increasing concentrations of NaBH_4_.

### MS analysis of NaBH_4_-treated LarA_Lp_

We next conducted MS experiments to understand how the enzyme is permanently inactivated by NaBH_4_. Because significant spectral changes were noted when using the reagent at 0.1 mM concentration and most of the changes were complete using 1 mM reductant, we chose three concentrations of NaBH_4_ (50, 100, and 250 μM) to treat the enzyme, and compared the results with the untreated sample (we included 2-3 replicates for each condition with representative spectra shown in **Figure 3A**). The untreated LarA_*Lp*_ sample contained a major species with a mass of 47845 Da, which is consistent with the expected mass of StrepII tagged holoenzyme tethered to an unmodified NPN cofactor. Addition of 50 μM NaBH_4_ to LarA_*Lp*_ drastically changed the MS spectrum, resulting in a major species with a mass of 47789 Da, a change of -56 Da (Δ - 56 Da) when compared to the holoenzyme. This species also is the major component for the samples treated with greater concentrations of NaBH_4_. One likely explanation for the formation of this species is that Ni is lost (−58.7 Da) while three (or two) hydrogen atoms are added to the protein. To compare the Ni content before and after NaBH_4_ treatment, both the untreated and treated samples were subjected to size-exclusion chromatography to replace the buffer, and the proteins were analyzed by inductively coupled plasma-mass spectrometry (ICP-MS). As shown in **Figure 3B**, the Ni/protein molar ratio in the sample treated with 2 mM NaBH_4_ was significantly decreased by more than 60% when compared to the untreated sample, supporting the notion that Ni dissociates from the enzyme after NaBH_4_ treatment.

**Figure 3.**
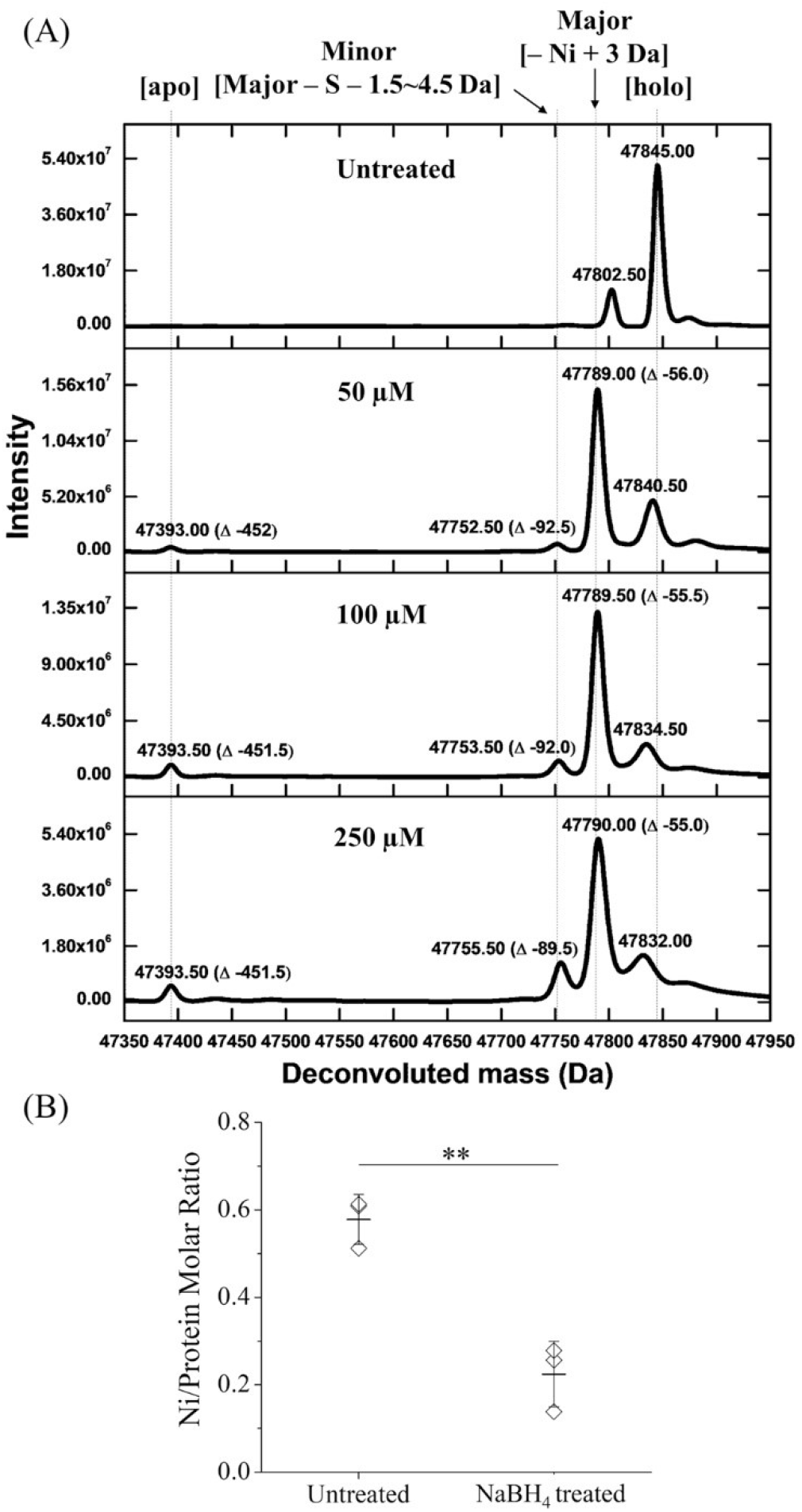
MS and Ni content changes of LarA_*Lp*_ caused by treatment with NaBH_4_. (**A**) MS analysis of LarA_*Lp*_ before and after treatment with the indicated concentrations of NaBH_4_. The monomeric LarA_*Lp*_ (∼20 μM) collected by size-exclusion chromatography in a buffer containing 50 mM Tris-HCl, pH 7.5, 125 mM NaCl, and 50 μM Na_2_SO_3_ was treated with NaBH_4_ for 10 min at room temperature. (**B**) Ni content analysis of LarA_*Lp*_ before and after treatment with NaBH_4_. The monomeric LarA_*Lp*_ (24-36 μM) collected as above was treated with 2 mM NaBH_4_ for 10 min at room temperature. After digestion with nitric acid, the Ni contents of the samples were measured using ICP-MS. The protein concentrations were determined based on the absorbance at 280 nm using a Nanodrop spectrophotometer. The results shown are from one representative experiment with three biological replicates. The asterisks indicate the significant differences (Student’s *t* test: ** P<0.01.)

Another noticeable change in the MS spectra of NaBH_4_-treated enzyme is the appearance of a feature corresponding to the apoenzyme (47393 Da, Δ -452 Da) for which the signal intensity relative to the major species increased as higher concentrations of NaBH_4_ were applied (**Figure 3A**). It was reported that NaBH_4_ treatment may lead to thioamide bond cleavage or sulfur loss in small molecules.^12^ Accordingly, it is conceivable that the thioamide bond linking the NPN cofactor to the lysine residue (K184) was partially disrupted by NaBH_4_ treatment, leaving a portion of the enzyme in the apoenzyme state. The reaction of the thioamide bond with hydride may also explain the formation of another minor species with a variable mass in the range of 47752.5-47755.5 Da (Δ -92.5∼-89.5 Da); this species may represent the loss of a sulfur atom from the Ni-free major species with additional elimination of an uncertain number of hydrogen atoms (in the range of 1-5, depending on the NaBH_4_ concentration).^12^ A scheme depicting these reactions involving the thioamide bond is shown in **Figure S3**. Because the signals associated with the apoprotein and the species missing a sulfur atom were much smaller than that of the major Ni-free species, the reactions leading to thioamide bond cleavage and sulfur loss are unlikely to be responsible for LarA_*Lp*_ inactivation.

Given that treatment with 250 μM NaBH_4_ nearly eliminated the LarA_*Lp*_ activity (**Figure 1**), the transition of holoenzyme to the major species of mass 47789 Da must solely account for the permanent inactivation. With the evidence supporting a hydride attack on C4 of the NPN cofactor (**Figure 2**), it is reasonable to believe that Ni loss triggered by hydride transfer completely abrogated activity. To better understand how the C-Ni bond is broken, we conducted a structural study on LarA_*Lp*_ with its NPN cofactor in the reduced state.

### Formation of the sulfite-NPN adduct at C4 weakens the C-Ni bond

Initial trials to crystallize the NaBH_4_-treated LarA_*Lp*_ with a reduced NPN cofactor failed, largely due to protein aggregation after NaBH_4_ treatment as shown by size-exclusion chromatography (**Figure S4**). A prior structural study had shown that sulfite can enter the active site as a substrate analog and form a covalent bond with C4 of pyridinium ring as a reducing agent.^9^ More generally, sulfite is known to form a covalent bond with hydride-accepting atoms in cofactors including NAD^+^ and FAD.^13^ Unfortunately, the presence of high concentrations of sulfate in the previously reported crystallization buffer (2 M ammonium sulfate) and the moderate resolution (2.4 Å) of the earlier structure had complicated interpretation of the density and compromised a more thorough structural characterization.

In an effort to solve the structure of LarA_*Lp*_ in the absence of sulfate, we generated a double variant (R98A/R100A) involving residues shown to interact with this anion and demonstrated that it was functional and crystallizable in the absence of sulfate or other multi-charged anions (**Figure S5**). The structure of this variant co-purified with sulfite was solved at 2.15 Å with an RMSD of 0.49 Å when compared with the closed state of LarA_*Lp*_ (PDB: 5HUQ, chain B), allowing us to revisit the sulfite-NPN adduct at improved resolution (**Table S1**). As shown in **Figure 4A**, the electron densities at the active site support a covalent modification of NPN by a ligand with a shape and size that are consistent with a sulfite molecule. Indeed, modeling the sulfite-NPN adduct (PDB ligand ID: ENJ) fits the density very well, and no other components in the buffers for purification or crystallization match the density as well as sulfite. To remove sulfite as much as possible to allow for co-crystallization with substrate, we replaced sulfite with 2 mM D-lactate during size-exclusion chromatography; however, the density remained consistent with sulfite, not lactate, in the structure that was solved at a resolution of 2.0 Å (**Figure 4B** and **Table S1**). This finding indicates that sulfite affinity for LarA_*Lp*_ is much greater than that for the natural substrate and the dissociation rate appears to be very low. Notably, modeling the sulfite-NPN adduct into this structure led to a negative density between sulfite and the NPN cofactor, whereas modeling a sulfite separated from the NPN cofactor resulted in an intervening positive density. This result indicates that the sulfite-NPN adduct becomes partially decomposed, consistent with the covalent bond between sulfite and the NPN cofactor undergoing slow cleavage at the active site. This ability to undergo bond breakage explains why the sulfite adduct was not observed in the MS spectra (**Figure 3**), why the enzyme co-purified with sulfite is functional, and why sulfite acts as a reversible inhibitor with a mixed inhibition mechanism ^3^.

**Figure 4.**
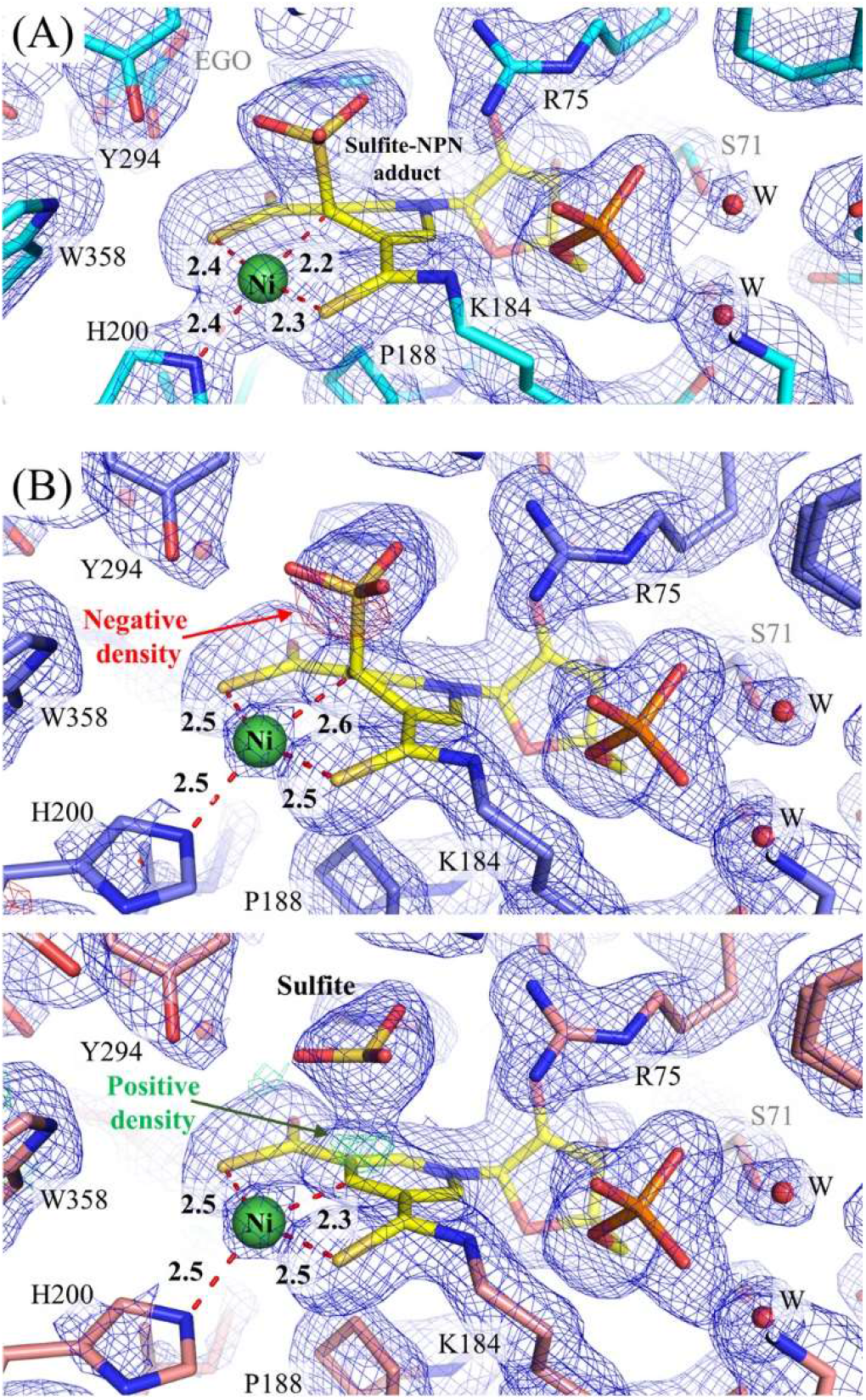
Crystal structures of the R98A/R100A variant of LarA_*Lp*_. (**A**) Sample that was co-purified with 50 μM sulfite. (**B**) Two models that were used to fit the data for sample initially purified with sulfite which was replaced with 2 mM D-lactate. *Upper panel*: Built with the sulfite-NPN adduct resulting in negative density at the covalent attachment site (mF_o_-DF_c_ map, σ = 3, red meshes). *Lower panel*: Built as separate sulfite and NPN cofactor resulting in positive intervening density (mF_o_-DF_c_ map, σ = 3, green meshes). 2mF_o_-DF_c_ maps (σ = 1) are depicted as blue meshes. The bond lengths around the Ni atom are labeled in angstroms. The structures are shown in stick mode and the residues located in close proximity to the cofactor are labeled. Water molecules are drawn as red spheres. EGO: ethylene glycol which is a component in the crystallization solution.

When enzyme samples containing the sulfite-NPN adduct and the unmodified NPN cofactor (PDB ligand ID: 4EY) were structurally compared, two major differences were noticed. Firstly, the sulfite-reduced pyridine ring adopts a boat configuration with C4 departing from the plane to form a C-S bond. The nitrogen atom in the ring also is out of plane, ending up as a tertiary amine. However, the angles around this atom (116.7°, 114.9°, 119.1°) suggest that it is of a mixed *sp2*-*sp3* hybridization (**Figure S6**). The deviation from the ideal geometry of a tertiary amine is consistent with the instability of the sulfite-NPN adduct. Secondly, the C-Ni bond becomes longer upon adduct formation. In the unmodified NPN cofactor, the C-Ni bond length is within the range of 2.0-2.1 Å (PDBs: chain B of 5HUQ; chain B and chain C of 6C1W), which is slightly longer than those in small molecule SCS-type Ni pincer complexes (1.8-1.9 Å) ^14, 15, 16^.

Upon formation of the adduct, the C-Ni distance increased to 2.2 Å (**Figure 4A**) or 2.6 Å (**Figure 4B**, lower panel). In the latter case, the completely broken C-Ni bond likely accounts for the very low occupancy for Ni (∼20%).

### Implications for the PCHT mechanism

Given that hydride transfer is a key step of the PCHT mechanism, one can ask why a hydride derived from NaBH_4_ leads to Ni loss and enzyme inactivation when it attacks C4 whereas a hydride derived from substrate that also attacks C4 does not lead to activity loss. The combined biochemical and structural data allow us to propose a reaction mechanism for LarA_*Lp*_ inactivation by NaBH_4_ (**Figure 5**), which helps to answer this question. In this mechanism, hydride attacks C4 of NPN, leading to pyridinium reduction and formation of the boat configuration where the C-Ni bond is weakened. Next, two possible paths lead to Ni loss. The Ni^2+^ may dissociate to create a C4 carbanion, and a proton from solvent can associate (Path A, **Figure 5**). In addition, an alternative path is based on the reactivity of a small molecule Ni pincer complex that, when reduced at C4, becomes unstable, resulting in irreversible demetallation due to rearomatization.^14^ If this occurs in LarA_*Lp*_ (Path B), the resulting pyridinium after Ni loss would be similar to the counterpart in NAD^+^, which may be then reduced by a second hydride from NaBH_4_. Through either path, the enzyme containing the reduced and Ni-free cofactor likely accounts for the permanent inactivation by NaBH_4_. Consistent with this notion, this species has been proposed to be in a stable state according to a computational study.^11^ When it comes to the racemization reaction, the substrate-derived hydride is transferred to C4 and then rapidly attacks the intermediate held at the active site. As a result, NPN is only transiently reduced with a probably very short lifetime. In contrast, the lack of a hydride acceptor at the active site in the NaBH_4_-involved reaction may increase the chance of further reactivity involving C4 such as the irreversible loss of Ni and addition of a proton from solvent. Indeed, water molecules are found in close proximity to the active site, particularly through a water channel connecting to the exterior of the enzyme (**Figure S7**). When the intermediate is positioned close to the reduced NPN cofactor, it may serve as a cap to insulate C4 against solvent and thus prevent C4 from unwanted reactivity involving Ni loss. Accordingly, by acting as both a hydride acceptor and a water-proof cap, the intermediate is postulated to play an active role in protecting the active site from the fatal side reaction(s) involving unprotected reduction of the NPN cofactor. A trapped intermediate in the active site chamber during the racemization reaction has been implicated in an early study of lactate racemase from *Clostridium butylicum*, in which ^14^C labeled pyruvate did not exchange with the unlabeled intermediate generated at the active site to yield a product with radioactivity.^17^ Our analysis also highlights a key difference between the LarA_*Lp*_ catalyzed reaction and those mediated by small molecule NPN mimics.^14, 16^ While the intermediate generated during the enzyme catalyzed reaction is retained in the active site, there is no mechanism for the small molecule NPN mimics to use the intermediate to protect the vulnerable reaction center. As a result, the small molecule NPN mimics are either unstable after hydride transfer ^14^ or millions of times slower than LarA_*Lp*_ in catalyzing racemization.^16^ Holding the intermediate at the reaction center with a proper orientation would be an effective strategy to improve catalysis efficiency and stability of the catalyst.

**Figure 5.**
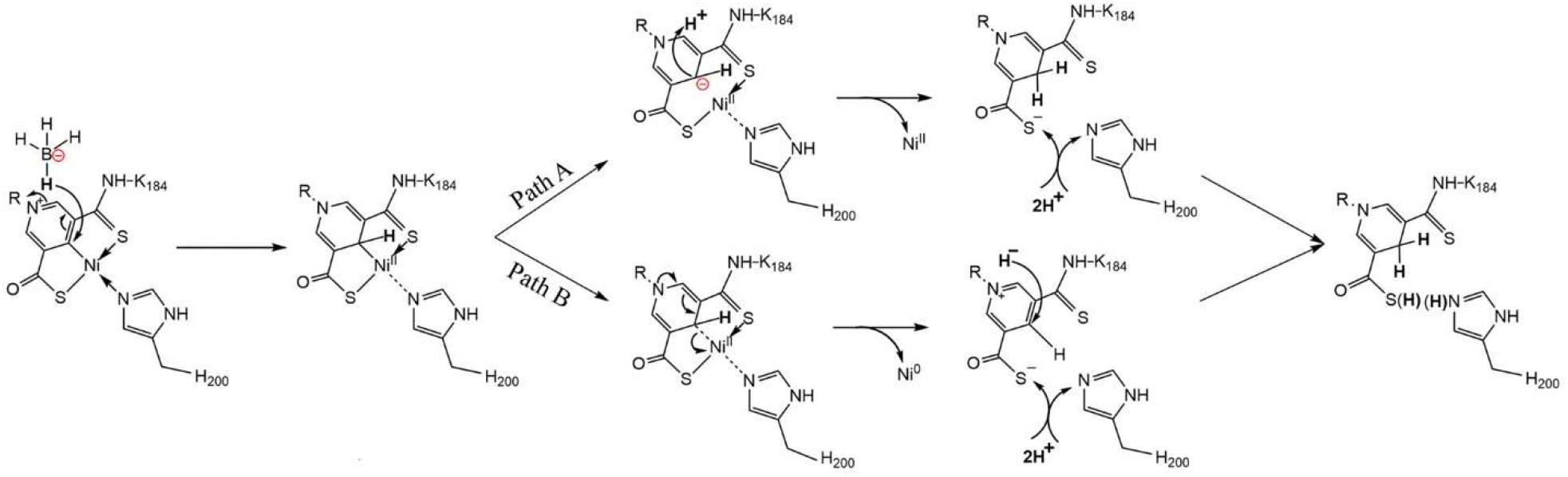
Proposed reaction mechanism for inactivation of LarA_*Lp*_ by NaBH_4_ involving chemical changes to the NPN cofactor. The hydride attack on C4 of the NPN cofactor leads to a weakened C-Ni bond. In Path A, dissociation of the Ni(II) from the cofactor results in carbanion formation at C4 which then acquires a proton from solvent. In Path B, the NPN cofactor undergoes rearomatization, loss of Ni(0), and a second hydride transfer to C4. The two paths converge at the reduced and Ni-free cofactor. These changes, together with partial protonation of the metal-free thioacid and His200, account for the molecular weight of the major species observed in MS experiments conducted in positive mode (Figure 3).

## Conclusion

In this work, our biochemical, spectroscopic, and MS analysis of LarA_*Lp*_ inactivation by NaBH_4_ consistently supported hydride transfer to C4 of the NPN cofactor. Different from the lactate racemization reaction in which the NPN cofactor is transiently reduced in the hidden redox reaction, hydride transfer to C4 during the NaBH_4_ treatment is followed by C4 reactivity involving irreversible Ni loss, which accounts for the permanent inactivation of the enzyme. As revealed in the high-resolution structures, the weakened C-Ni bond upon the formation of the sulfite-NPN adduct provides the structural basis for this vulnerability of the C-Ni bond when the NPN cofactor is in the reduced state. Collectively, these findings add new structural and biochemical insights into the PCHT mechanism for LarA_*Lp*_, which may shed light on how to design more potent small molecule NPN mimetics for potential applications.

## Experimental procedures

### Protein expression and purification

LarA_*Lp*_ was produced in *Lactococcus lactis* expression strain NZ3900 cells bearing the plasmid pGIR112 as previously described.^3^ This plasmid contains *larA*, fused at the 5’ end to a DNA sequence encoding a StrepII tag, as well as *larB, larC*, and *larE* that encode the biosynthetic enzymes to generate the NPN cofactor. Site-directed mutagenesis was conducted by using the QuikChange® kit to generate the R98A/R100A double variant (**Table S2**) that was expressed in the same cells. The cell pellets were stored at -80°C until further use. The cell pellets were thawed, resuspended in 80 ml of lysis buffer (100 mM Tris-HCl, pH 7.5, 150 mM NaCl, 50 μM Na_2_SO_3_, 10 μg/ml lysozyme and 2 μg/ml of DNaseI), and stirred for 1 h at 4°C. Phenylmethylsulfonyl fluoride (1 mM) was added to the resuspension and cell lysis was performed by sonicating the samples with cycles of 5-s on/10-s off on ice for 15-20 min. The cell lysates were centrifuged at 18,500 *g* for 1 h and the clarified supernatant solutions were loaded onto 2 ml of Streptactin-XT resin with 1 min/ml flow rate. For each sample, the resin was washed with 8 column volumes (CV) of wash buffer (100 mM Tris-HCl, pH 7.5, 150 mM NaCl, and 50 μM Na_2_SO_3_) and eluted with 20 CV of elution buffer (wash buffer plus 50 mM biotin). The samples were concentrated and loaded onto Superdex 200 increase 10/30 GL columns that were equilibrated with the gel filtration buffer (50 mM Tris-HCl, pH 7.5, 125 mM NaCl, and 50 μM Na_2_SO_3_). To purify the proteins in the absence of sulfite, Na_2_SO_3_ in the gel filtration buffer was replaced with 2 mM D-lactate.

### Lactate racemase assay and inhibition of LarA_Lp_ by NaBH_4_

1 M NaBH_4_ was freshly prepared in 1 M NaOH and diluted to 100 mM in 50 mM NaOH as the stock solution and used immediately after preparation. The dose dependent inactivation (NaBH_4_, 5 μM-2 mM) of LarA_*Lp*_ (∼20 μM) was performed in the buffer containing 50 mM Tris-HCl, pH 7.5, 125 mM NaCl, and ∼0.1 μM Na_2_SO_3_, and the activity of the NaBH_4_ treated enzyme (340 ng for NaBH_4_ inactivation, or 200 ng for comparison of the R98A/R100A variant with the wild-type enzyme) were conducted in the reaction buffer containing 60 mM MES, pH 6.0, and 20 mM D-lactate. The reactions were started by adding the enzyme to the reaction buffer, followed by incubation at 35°C for 10 min and termination by heating at 95°C for 10 min. The precipitated protein was pelleted by centrifugation and the L-lactate concentration was measured using the D-/L-lactate assay kit (Megazyme Inc.), as previously described.^3^ To examine the reversibility of inactivation, purified LarA_*Lp*_ was treated with 2 mM NaBH_4_ and desalted by ultrafiltration using a concentrator (Amicon ultra, 30 kDa) and a buffer containing 50 mM Tris-HCl, pH 7.5, and 125 mM NaCl.

### UV-Vis spectroscopy

The monomeric species of LarA_*Lp*_ (∼20 μM) from the gel filtration step (50 mM Tris-HCl, pH 7.5, 125 mM NaCl, and 50 μM Na_2_SO_3_) was treated with NaBH_4_ (0.1-2.5 mM) and incubated for 10-15 min at room temperature until the bubbles (hydrogen gas) dissipated. The UV-Vis absorption spectra (300-700 nm) were recorded for the untreated and NaBH_4_ treated samples on a Shimadzu UV-2600 spectrophotometer at 25°C.

### Protein MS experiment

The monomeric species of LarA_*Lp*_ from the gel filtration step was used immediately after purification to perform NaBH_4_ treatment and MS analysis. Purified protein (∼20 μM) was treated with NaBH_4_ at 50 μM, 100 μM, or 250 μM at room temperature for 10 min. All samples were transferred to 4°C and centrifuged at 12,000 rpm for 10 min. Each sample was diluted to 1:40 with 50 mM Tris-HCl, pH 7.5, and 125 mM NaCl to perform MS analysis. Samples (10 μl) of untreated and NaBH_4_-treated LarA_*Lp*_ protein solutions were injected onto a cyanopropyl column connected to Xevo G2-XS QTof (Waters) with a flow rate of 0.1 ml/min. The chromatography used 0.1% formic acid (solvent A) and acetonitrile (solvent B) as solvents in a 98%/2% ratio. Scanning was performed at 1 scan/s in positive ion mode.

### ICP-MS experiment

The monomeric species of LarA_*Lp*_ purified from three separate batches of size-exclusion chromatography was used immediately for performing the NaBH_4_ treatment. The protein (24-36 μM) was treated with 2 mM NaBH_4_ at room temperature for 10 min. Both treated and untreated samples from the same batch were applied to size-exclusion chromatography to remove Ni dissociated from the holoenzyme. The protein-containing fractions were pooled and concentrated to 12-26 μM. The samples were digested with 70% nitric acid and then diluted by 30 times before loaded to Agilent 8900 Triple Quadrupole ICP-MS to determine Ni content.

### Crystallization, data collection, processing, and structure determination

The concentrated R98A/R100A variant protein (20 mg/ml) was screened with the commercially available kits (Index, JCSG^+^, Crystal screen, and Morpheus). For the protein purified in the presence of sulfite, plate shaped crystals were obtained in 100 mM sodium HEPES/MOPS, pH 7.5, 60 mM MgCl_2_·6 H_2_O, 60 mM CaCl_2_·2 H_2_O, 20% ethylene glycol and 10% PEG 8000. In the case of protein purified with 2 mM D-lactate, the best diffracting crystals were obtained with 27% ethylene glycol and 14% PEG 8000 as precipitant in the same buffer. The crystals were flash frozen in liquid nitrogen.

The diffraction data of the crystals were collected at the Life Sciences Collaborative Access Team (LS-CAT) beamline 21-ID-G at the Advanced Photon Source in the Argonne National Laboratory, and processed with HKL2000.^18^ The structures were solved by molecular replacement in Phenix.phaser using the wild-type LarA_*Lp*_ structure (PDB: 5HUQ, chain B) as the search model, and refined with Phenix.refine.^19^ Model building was conducted in Coot.^20^ The 2mF_o_-DF_c_ and mF_o_-DF_c_ maps were generated in Phenix and the protein structure figures were prepared in Pymol v 2.4.1 (Schrodinger). The atomic coordinates have been deposited in the PDB with accession IDs 8EZF, 8EZH and 8EZI.

## Acknowledgements

We thank Dr. Tony Schilmiller in the Mass Spectrometry & Metabolomics Core Facility for helping in the protein mass spectrometry experiments. We thank Dr. Keith MacRenaris in the Quantitative Bio Element Analysis and Mapping Facility and Department of Microbiology and Molecular Genetics and Yuhan Jiang in Department of Biochemistry & Molecular Biology for helping in the ICP-MS experiments. We thank the beamline scientists of LS-CAT at APS for assisting data collection. We thank Kevin Kamil Jumaa for assisting with regular lab experiments.

## Author Contribution

J.H. and R.P.H. formulated the project. S.G. and D.S. conducted experiments. S.G., R.P.H. and J.H. analyzed data and wrote the manuscript.

---

**Scheme 1.**
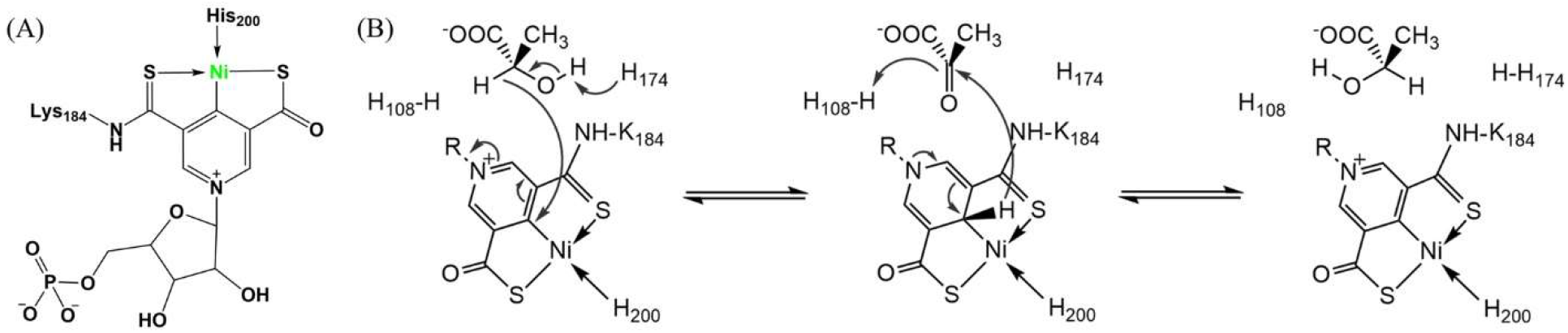
Structure of the NPN cofactor **(A)** and racemization of D-/L-lactic acid by LarA_*Lp*_ via a proton-coupled hydride-transfer (PCHT) mechanism **(B)**.

