## Supplementary information for "Irreversible inactivation of lactate racemase by sodium borohydride reveals reactivity of the nickel-pincer nucleotide cofactor"

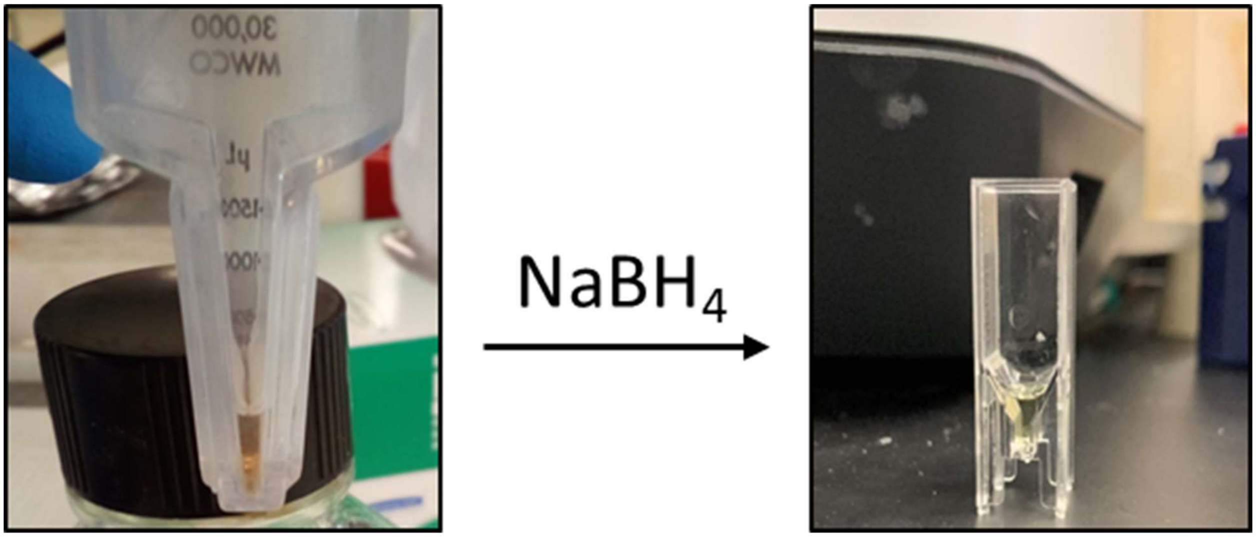


**Figure S1.** Color change of purified $LarA_{Lp}$ upon $NaBH_4$ treatment. The sample depicted on the left is $LarA_{Lp}$ (116 μM) that was purified in the presence of 50 μM $Na_2SO_3$ in a buffer containing 100 mM Tris-HCl, pH 7.5, and 125 mM NaCl. On the right is a sample (20 μM) purified in the same way and treated with 2 mM $NaBH_4$.

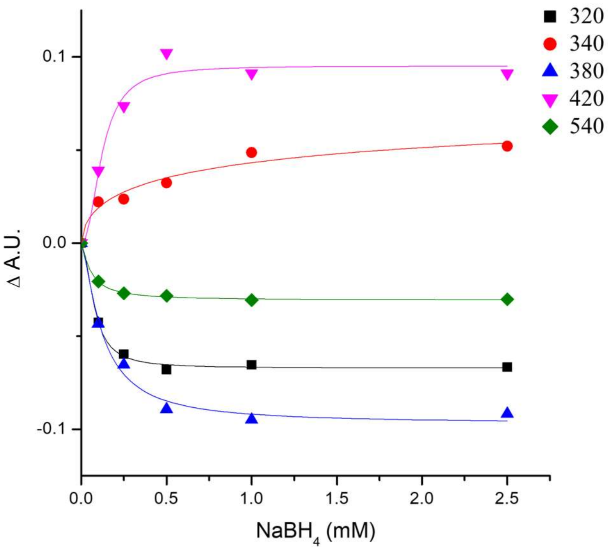


$$y = A \cdot x^n \; / \; (K^n + x^n)$$

| Wavelength (nm) | K (mM) | n | $r^2$ |
|---|---|---|---|
| 320 | 0.075 ± 0.0068 | 1.9 ± 0.42 | 0.996 |
| 340 | 0.95 ± 3.1 | 0.56 ± 0.4 | 0.94 |
| 380 | 0.12 ± 0.018 | 1.4 ± 0.36 | 0.98 |
| 420 | 0.12 ± 0.019 | 2.2 ± 0.79 | 0.96 |
| 540 | 0.056 ± 0.0066 | 1.3 ± 0.23 | 0.998 |

**Figure S2.** UV-Vis absorption changes at the indicated wavelengths during the titration of LarA*Lp* with $NaBH_4$. The dose-dependent curves were fitted using the Hill model and the results are shown in the table. K represents the apparent affinity, n is the Hill coefficient, and r is the correlation coefficient generated in curve fitting. Note that the quality of curve fitting of the absorption changes at 340 nm is worse than the others, likely due to the interference by the greater spectral changes at 320 nm and 380 nm.

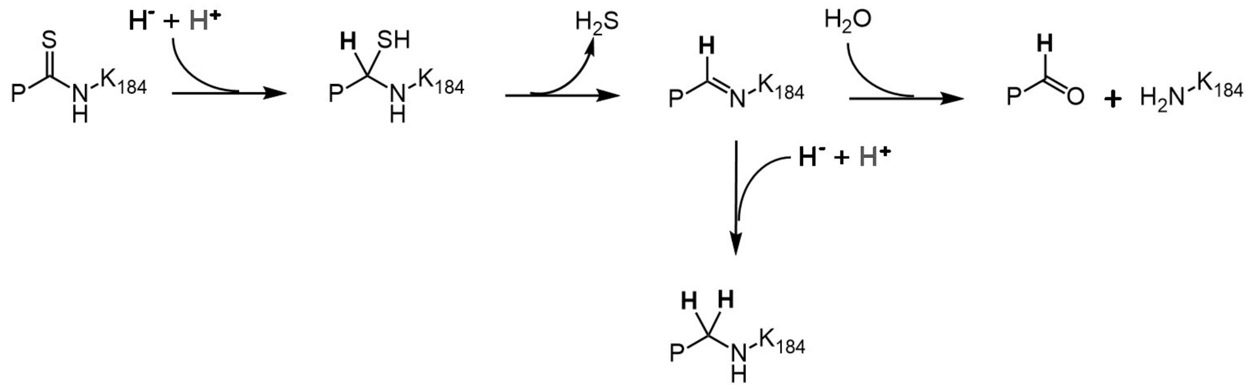


**Figure S3.** Proposed reactions involving the thioamide bond upon $NaBH_4$ treatment. The thioamide bond, which links the NPN cofactor and K184 of LarA$_{Lp}$, is attacked by hydride, followed by hydrogen sulfide elimination to form a Schiff base, which may undergo hydrolytic cleavage to generate the apoenzyme, or be further reduced by hydride to generate the species missing one sulfur atom. P: pyridinium of the NPN cofactor.

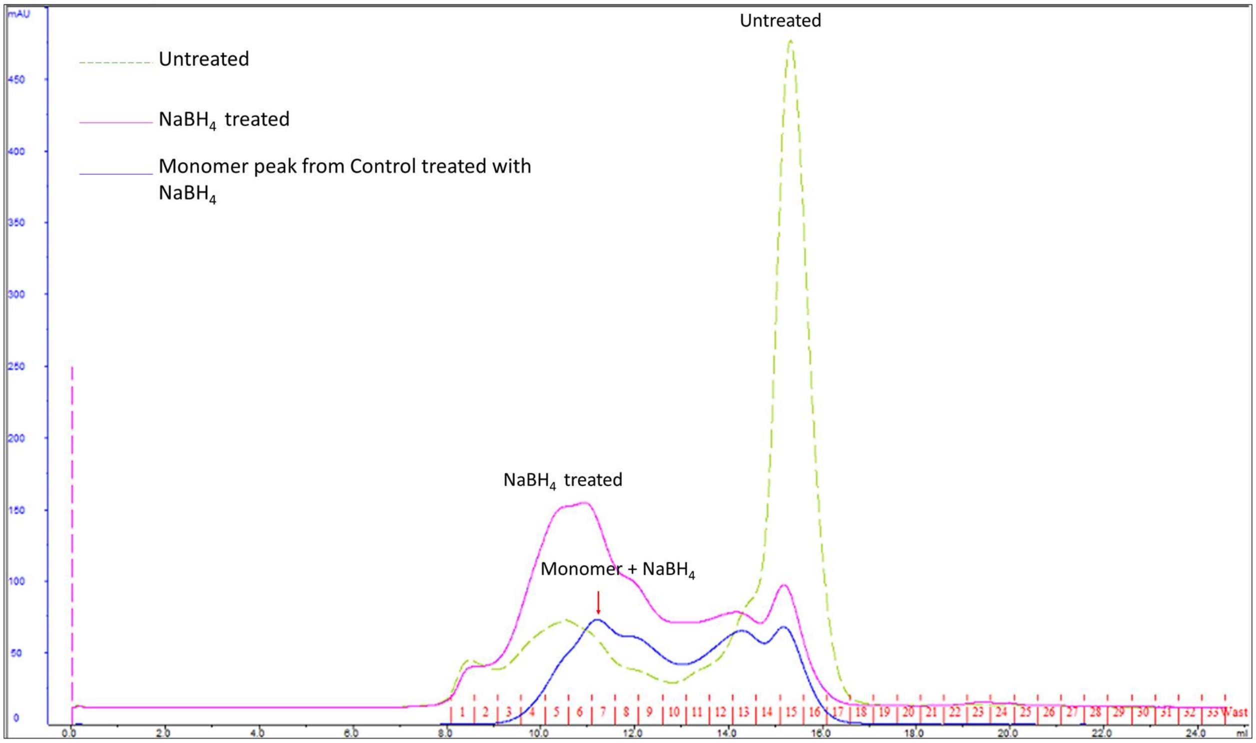


**Figure S4.** Size-exclusion chromatography profiles of the $NaBH_4$-treated R98A/R100A variant of LarA$_{Lp}$ in comparison with the untreated sample. Samples were chromatographed on a Superdex 200 increase 10/30 GL column equilibrated with a buffer containing 50 mM Tris-HCl, pH 7.5, and 125 mM NaCl. The control sample (green dashed line) is the untreated variant, the "$NaBH_4$ treated" sample (pink solid line) is the variant protein purified by affinity chromatography and then treated with 0.5 mM $NaBH_4$, whereas the "Monomer + $NaBH_4$" sample (blue solid line) is the monomeric fraction collected from the untreated sample which was then treated with 0.5 mM $NaBH_4$. $NaBH_4$ treatment led to protein aggregation.

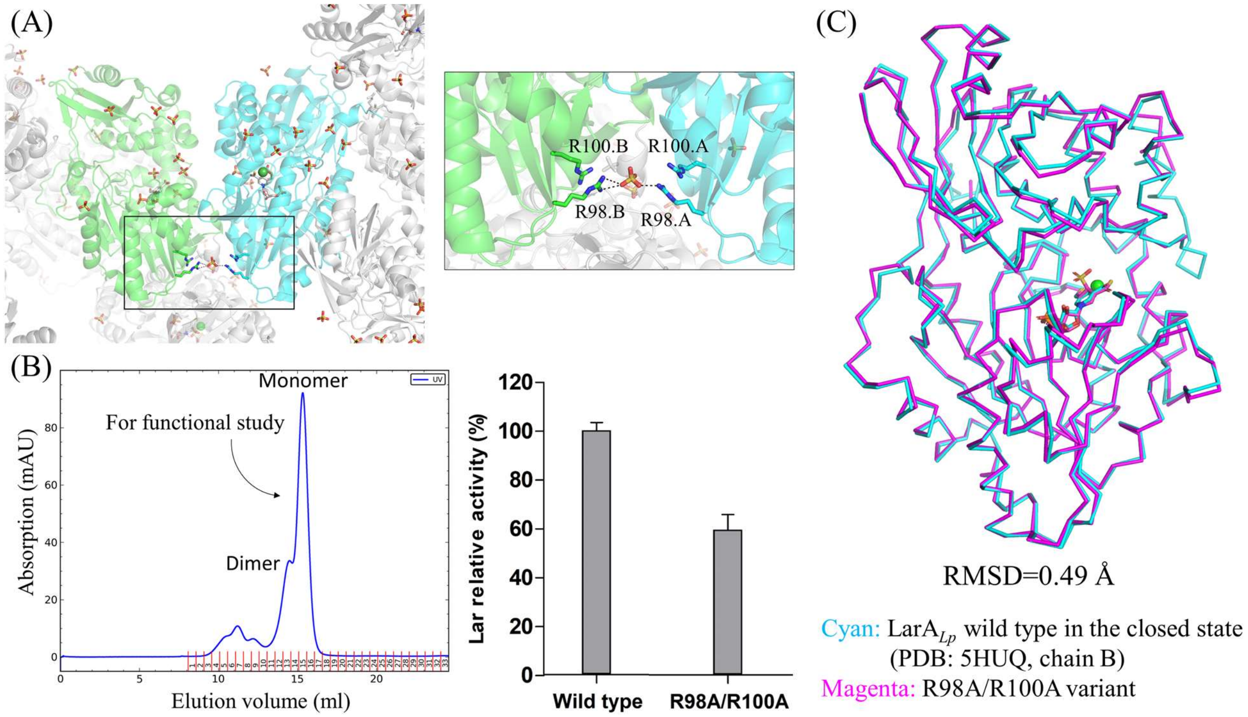


**Figure S5.** Design, characterization, and structure determination of the R98A/R100A double variant of LarA$_{Lp}$. (**A**) Crystal packing analysis of the wild-type LarA$_{Lp}$ identified a sulfate binding site at an arginine cluster located at the crystallographic dimerization interface. Elimination of this sulfate binding site reduced the dependence of crystallization on multi-charged anions. (**B**) Purification and lactate racemization activity assay of the double variant in comparison with the wild-type enzyme. 200 ng of enzyme was used in this assay. (**C**) Structural comparison of the double variant with the wild-type enzyme. The overall RMSD is 0.49 Å.

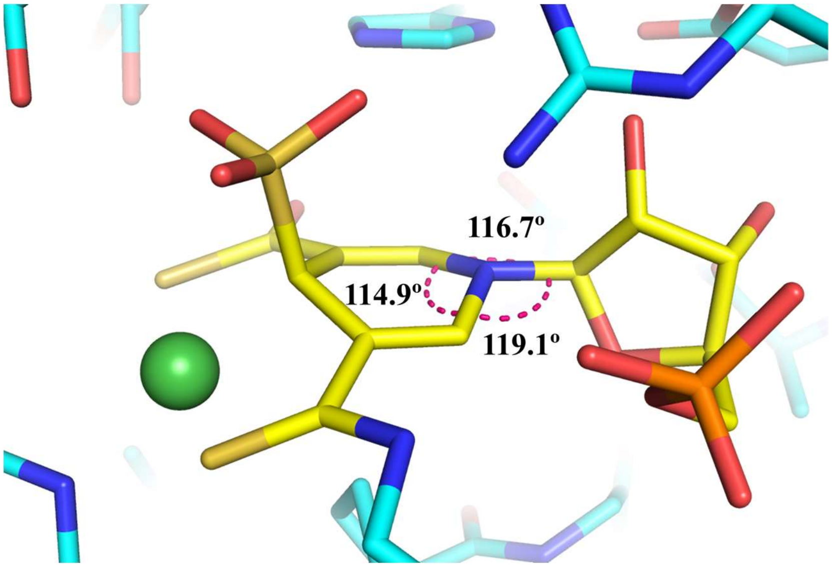


**Figure S6.** Angles around the N atom of the pyridine in the sulfite-reduced NPN cofactor. The electron density map of the same structure is shown in Figure 4A.

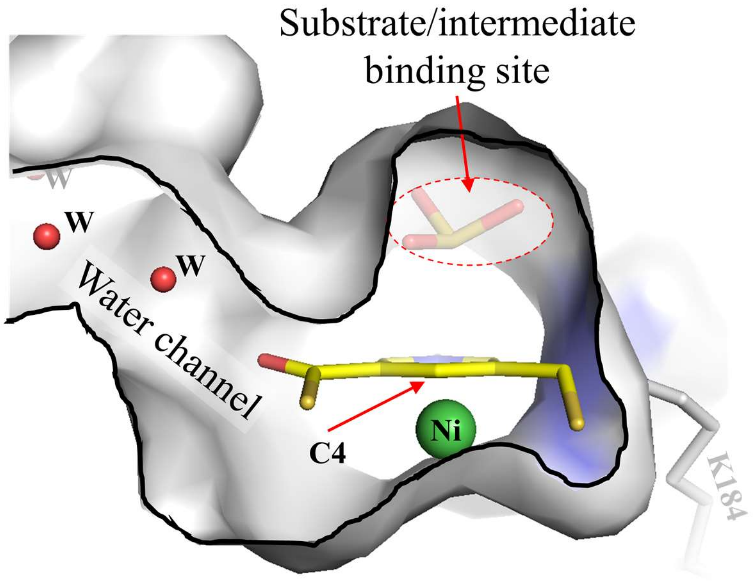


**Figure S7.** The active site of LarA$_{Lp}$ with bound sulfite shown in surface mode (grey). The cofactor (yellow sticks) and the substrate analog sulfite are depicted in the cavity of the active site with a water channel connecting to the exterior of the protein. The pyruvate intermediate that is formed *in situ* appears to insulate C4 against solvent. We postulate that the intermediate may play a role in protecting C4 from fatal side reaction(s) involving unprotected reduction of the NPN cofactor.

**Table S1.** Crystallographic statistics

| **Data collection** | Co-purified with sulfite | Purified initially with sulfite and then replaced with 2 mM lactate | |
|---|---|---|---|
| Beamline | 21-ID-G | 21-ID-G | |
| Wavelength (Å) | 0.97856 | 0.97856 | |
| Space group | $P\ 2_1\ 2_1\ 2_1$ | $P\ 2_1\ 2_1\ 2_1$ | |
| Unit cell a, b, c (Å); α, β, γ (°) | 42.84 ,79.91, 121.78<br>90, 90, 90 | 43.01, 79.09, 121.17<br>90, 90, 90 | |
| [a]Resolution (Å) | 37.97 – 2.15 (2.22– 2.15) | 37.79 – 1.99 (2.06-1.99) | |
| Unique reflections | 23,345 (1,921) | 29,029 (2074) | |
| [a]Redundancy | 7.8 (6.9) | 10.0 (9.1) | |
| [a]Completeness (%) | 100 (99.8) | 100 (100) | |
| [a]$I/\sigma I$ | 10.0 (1.4) | 9.1 (1.3) | |
| [a,b]$R_{merge}$ | 0.21 (1.1) | 0.74 (4.7) | |
| [a,c]$R_{pim}$ | 0.079 (0.45) | 0.24 (1.62) | |
| [d]$CC_{1/2}$ | 0.65 | 0.42 | |
| **Refinement** | | Modeled with sulfite-NPN adduct | Modeled with separated sulfite and NPN |
| Protein atoms | 3153 | 3170 | 3170 |
| $H_2O$ molecules | 175 | 209 | 209 |
| Cl atoms | 1 | 3 | 3 |
| Ni atom | 1 | 1 | 1 |
| Calcium | 2 | 2 | 2 |
| Magnesium | 2 | 2 | 2 |
| $SO_3$ molecules | 0 | 0 | 1 |
| Ethylene glycol molecules | 52 | 7 | 7 |
| ENJ: reduced P2TMN<br>4EY: P2TMN | 1<br>0 | 1<br>0 | 0<br>1 |
| [e]$R_{work}/R_{free}$ | 0.1864/0.2430 | 0.1826/0.2213 | 0.1825/0.2211 |
| $B$-factors (Å$^2$) | 27.6 | 25.0 | 25.0 |
| Protein atoms | 27.3 | 24.8 | 24.7 |
| $H_2O$ molecules | 34.0 | 28.5 | 28.4 |
| Cl atoms | 48.7 | 43.0 | 42.9 |
| Ni atom | 39.0 | 33.1 | 34.4 |
| Calcium | 21.1 | 18.6 | 18.3 |
| Magnesium | 38.0 | 22.6 | 22.6 |
| Sulfite | 0 | 0 | 42.5 |
| Ethylene glycol | 29.9 | 26.7 | 26.7 |
| ENJ: reduced P2TMN<br>4EY: P2TMN | 0<br>21.8 | 26.3<br>0 | 0<br>24.5 |
| R.m.s. deviation in bond lengths (Å) | 0.008 | 0.008 | 0.008 |
| R.m.s. deviation in bond angles (°) | 0.972 | 1.04 | 1.01 |
| Ramachandran plot (%) favored | 97.6 | 96.6 | 96.6 |
| Ramachandran plot (%) outliers | 0 | 0 | 0 |
| Rotamer outliers (%) | 0.89 | 1.19 | 1.19 |
| PDB ID | 8EZF | 8EZH | 8EZI |

[a]Highest resolution shell is shown in parentheses.

[b]$R_{merge} = \sum_{hkl} \sum_j |I_j(hkl) - \langle I(hkl)\rangle| / \sum_{hkl} \sum_j I_j(hkl)$, where $I$ is the intensity of reflection.

[c]$R_{pim} = \sum_{hkl} [1/(N-1)]^{1/2} \sum_j |I_j(hkl) - \langle I(hkl)\rangle| / \sum_{hkl} \sum_j I_j(hkl)$, where N is the redundancy of the dataset.

[d]$CC_{1/2}$ is the correlation coefficient of the half datasets.
[e]$R_{work} = \sum_{hkl} | |F_{obs}| - |F_{calc}| | / \sum_{hkl} |F_{obs}|$, where $F_{obs}$ and $F_{calc}$ are the observed and the calculated structure factors, respectively. $R_{free}$ is the cross-validation R factor for the test set of reflections (5% of the total) omitted in model refinement.

**Table S2.** Strains, plasmids, and primers used in this study

| Strain, plasmid, or primer | Characteristics or sequences | Description or Source or reference |
|---|---|---|
| **Strains** | | |
| ***L. lactis* NZ3900** | MG1363 derivative | |
| **Plasmids** | | |
| **pGIR112** | Cmr; pNZ8048 with a 4.92-kb insert containing the translational fusion between PnisA and larA-larE with the Strep-tag fused at the C-terminus of LarA. | Ref2 |
| **pGIR112R98AR100A** | pGIR112 with the R98A/R100A mutation | This study |
| **Primers** | | |
| **R98AR100A-F** | CGG TCA GTG GCG CCC GAT GCA GCC ATT GCC ATC CTC GTA GCT ACT GGT TTC | |
| **R98AR100A-R** | GAA ACC AGT AGC TAC GAG GAT GGC AAT GGC TGC ATC GGG CGC CAC TGA CCG | |

References
